# Optimizing Cryo-FIB Lamellas for sub-5Å in situ Structural Biology

**DOI:** 10.1101/2022.06.16.496417

**Authors:** Sagar Khavnekar, Veronika Vrbovská, Magda Zaoralová, Ron Kelley, Florian Beck, Sven Klumpe, Abhay Kotecha, Jürgen Plitzko, Philipp S. Erdmann

## Abstract

We here present a method based on metallic platinum sputtering that can substantially enhance the quality of subtomogram averages from lamellas and thereby reduce the number of particles needed for high-resolution subtomogram averaging. We provide evidence for the physical background of this improvement and demonstrate its usefulness by producing sub-5Å ribosome averages from yeast.

## Main Text

Cryo-electron tomography (cryo-ET) is evolving into the method of choice for elucidating biological structures in their native environment. Together with subtomogram averaging (STA), it offers a unique way of imaging biological complexes in a near to native state and at subnanometer resolution. In recent years, advances in microscope hardware, data collection, and computational algorithms ^1–4^ have facilitated not only resolving various macromolecules in purified samples and whole cells at better than 5 Å resolution ^5,6^, but also directly reveal how different conformational states of complexes are linked to biological function. However, most model systems, such as *Saccharomyces cerevisiae* (*S. cerevisiae*), *Chlamydomonas reinhardtii* (*C. reinhardtii*), or *Caenorhabdidis elegans* (*C. elegans*) are too large to be imaged using transmission electron microscopy directly. In such cases, cryo-focused ion-beam (FIB) milling has become a widespread technique ^7–10^. However, averages from cryo-FIB samples which reach sub-5 Å resolution are still the exception and confined to high symmetry proteins or virus particles^11,12^.

This discrepancy between the achievable resolution of purified macromolecules as well as thin whole cells, and that from FIB-milled samples has remained unaddressed so far. In fact, many experimental parameters (e.g. sample thickness, defocus range, and number of targeted particles) are comparable between datasets from whole cells and lamellas. The few accounts of sub nanometer resolution averages from FIB-milled samples suggest that there is no fundamental barrier. Neither the damage layer nor the ion implantation by focused ion beam milling, which should significantly influence high resolution averaging^12,13^. It therefore stands to reason that either sample behavior in the microscope or imaging physics could be different for *in vitro* and *in situ* samples. Among many factors, beam-induced sample movement (BIM) and charging have long been discussed as confounding for high-resolution cryo-electron microscopy (cryo-EM). Charging has been found to be detrimental when imaging thicker samples (such as cellular lamellas) with the Volta phase plate (VPP)^15^. It is therefore common to sputter-coat cryo-FIB lamellas with a thin conductive platinum layer to mitigate the effects of specimen charging^16^ during VPP imaging. This concept has also been discussed in the context of micro electron diffraction to improve data quality^17^. In contrast, post milling sputter coating for defocus-based imaging has not been explored, most likely due to the granularity of the platinum layer (Supplementary Figure 1A) and its potentially negative effect on tomogram quality (Supplementary Figure 1B). While methods exist to computationally remove the sputtered platinum during reconstruction,^18^ it has not yet been adopted for high resolution subtomogram averaging.

To assess the effect on the achievable STA resolution, we compared samples with and without metallic Platinum coating. First, sputter conditions had to be improved to minimize image quality degradation by the Pt-layer. To obtain a fine and uniform conductive layer, the granularity of the platinum particles had to be considered and optimized (Supplementary Figure S1 A-B; detailed instructions in the Methods section). This is also required due to the subtle differences found between sputter coaters used on the various FIB-SEM tools.

With optimized conditions, including time, pressure and voltage, that avoided significant reconstruction artifacts, two cryo-ET datasets (comprising a total of 118 tilt series) were recorded to investigate the influence of Pt-coating on the final data quality. In brief, *S. cervisiae* cells were plunge frozen on carbon support EM grids and lamellas were automatically milled, (see Methods for details)^19^. On one set, metallic platinum was deposited after milling (+Pt) with the integrated magnetron plasma coater, while the other was left uncoated (−Pt). Tomograms were then recorded on a Krios G4 microscope with Selectris X energy filter (10 eV slit) and Falcon 4 detector using a dose symmetric tilt scheme (See Methods for details, Table 1). The +Pt dataset comprised 64 and the −Pt 54 tilt series. After template matching, subtomogram averaging and classification (see methods for details), the +Pt list contained ~12.5k, and −Pt ~9.5k particles in total. For comparison, the +Pt data was reduced to 9.5k particles by randomly selecting a subset to avoid any bias due to particle location, initial scoring, or defocus spread. While both datasets were matched in lamella thickness, defocus spread, particle positions and residual reconstruction errors (Supplementary Figure S2), the +Pt 80S ribosomes resulted in a final average at 5.1 Å global resolution (Supplementary Figure S3 A) while the −Pt dataset only reached 6.1 Å (Supplementary Figure S3 B). In both cases, local resolution extends to the resampled Nyquist frequency of 4 Å (Figure 1 A-B), but there is a two-fold increase in voxels with sub-4.5 Å resolution for +Pt (Supplementary Figure S3 C). To further assess the quality of the data, Rosenthal-Henderson B-factors^20^ were calculated for both sets. This revealed a significant reduction for the B-factor of sputter-coated cryo-FIB samples by 42% (Figure 1C). As both datasets have undergone the same sample preparation, data collection, computational processing and are matched in all characteristic parameters (see above), the improvement in the B_overall_ can be attributed entirely to the reduction in intrinsic amplitude decay of the images (B_image_) for the samples that were collected +Pt.^20^

**Figure 1.**
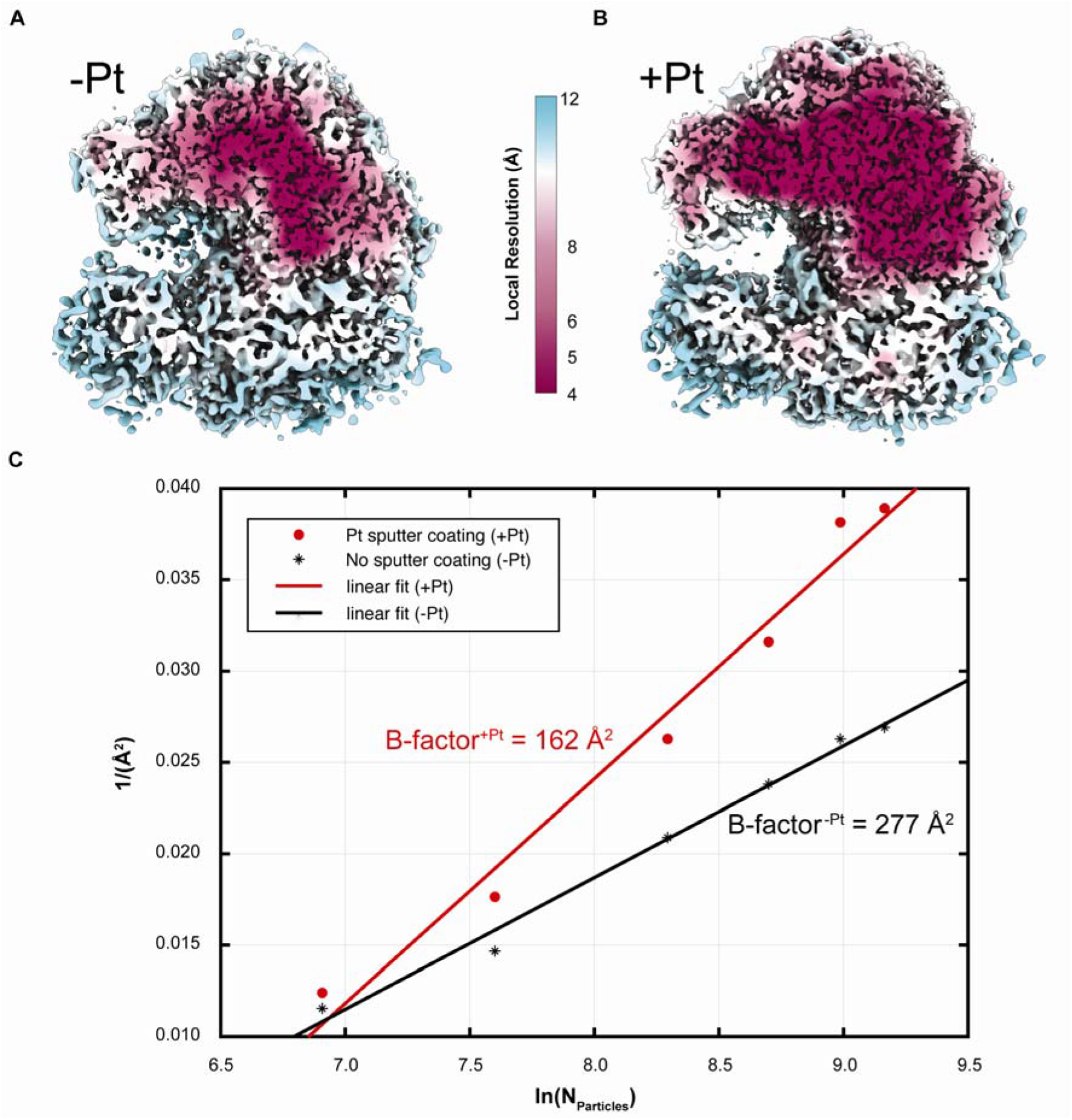
Sputter coating cryo-FIB milled lamellas with conductive platinum improves B-factors. A and B) Local resolution estimates mapped on cross-sections of the EM densities show a substantial increase from the −Pt (A) to the +Pt (B) dataset in pixels with resolution better than 4.5 Å. C) Rosenthal-Henderson B-factor plots reveal a significant improvement of the B-factor (42%) for the platinum-coated lamellas.

To elucidate where this improvement is coming from, local defocus estimates and sample movement were analyzed for each set. While changes in defocus did not differ significantly (Figure 2 A), mean accumulated shifts were on average five times larger for the −Pt than for the +Pt dataset (Figure 2 B). Additionally, local motion, as apparent from the median grid movement after Warp/M postprocessing^21^, is significantly larger for the sample lacking the conductive layer (Figure 2C). We therefore reason, that Pt-coating improves data quality mainly through a reduction in beam-induced sample movement (BIM). While BIM can be compensated for in tomography by motion correction, it has its limits due to the comparatively low dose and signal in individual movie frames. This may be an important difference to the single particle analysis (SPA) method, where each dose fraction receives a considerably higher electron dose and hence has better signal so that motion correction can be performed more efficiently. Reducing sample movement from the beginning, *i.e*. also within individual dose fractions, may therefore be more essential for *in situ* cryo-ET to retain the high-resolution information.

**Figure 2.**
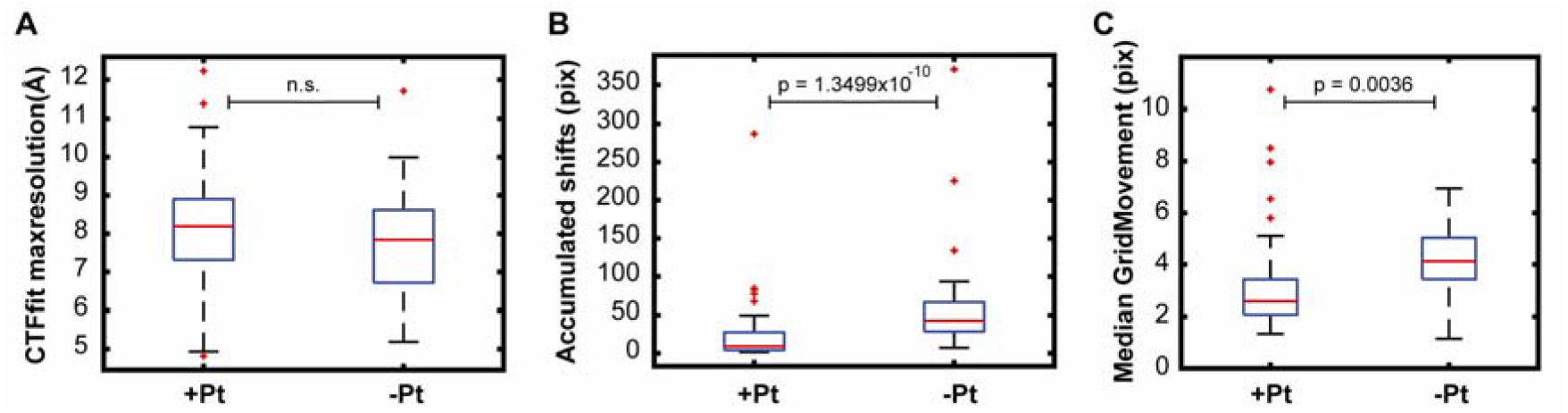
Global and local motion are significantly reduced for the +Pt dataset. A) CTF estimates (CTFfit max resolution) after the motion correction are not significantly different for +Pt and −Pt. B) Accumulated shifts are reduced by ~5x in the +Pt case. The medians are 8 (+Pt) vs. 42 pixels (−Pt). C) Local grid movements after tomographic tilt series refinement are also significantly lowerin the +Pt dataset (20%). On each box, the central mark indicates the median, and the bottom and top edges of the box indicate the 25th and 75th percentiles, respectively. The whiskers extend to the most extreme data points not considering outliers. Outliers are plotted individually using the ‘+’ marker symbol. P-values are calculated using the two-sample Kolmogorov-Smirnov test. n.s. = not significant.

To explore the resolution potential of the platinum-coated tomograms, the full +Pt dataset (91 tilt series) was subjected to the subtomogram averaging and classification pipeline. This yielded ~12.5k particles resulting in a map at 4.8 Å global resolution after 3D refinement. Subtomogram alignment focused on the 80S large subunit (LSU) and subsequent tilt-series refinement^5^ resulted in a final map at 4.5 Å global resolution. Local resolution, however, ranged between 3.5 - 4 Å for the LSU core (Figure 3A-B). The base stacking of the eukaryotic ribosomal RNA can clearly be resolved *in situ* at this resolution (Figure 3C). The high-resolution also facilitates identification and assignment of bulky side chains in alpha helices and beta sheets (Figure 3 D-E).

**Figure 3.**
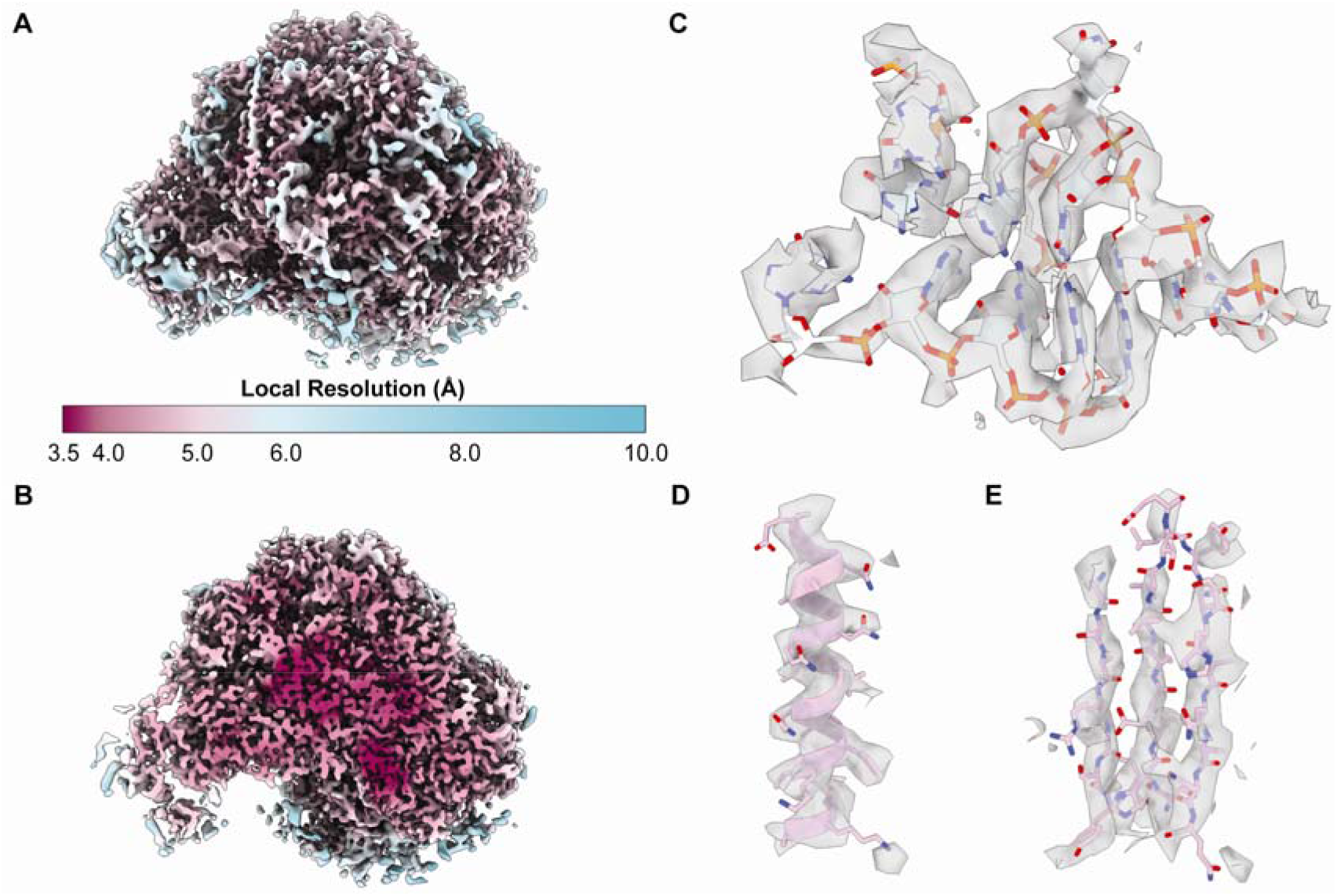
Local resolution differences and high-resolution features resolved in the +Pt dataset. A) Local resolution estimates mapped on the large ribosomal subunit (LSU). There is a significant improvement of voxels with a resolution of 4.5 Å or better for the +Pt map. B) Cross-section through the +Pt LSU core showing local resolution below 4 Å. C) RNA base stacking as well as and bulky amino acid side chains (D, E) are clearly resolved in these active ribosomes from lamellas.

In summary, mitigating sample charging is essential for reducing local beam-induced motion and obtaining high resolution subtomogram averages from cryo-FIB milled lamellas. This is demonstrated for one of the most abundant cellular complexes, the eukaryotic 80S ribosome. From a reasonable number of lamellas and tomograms, high resolution averages, which allow an unambiguous assignment of bulky amino acid side chains, as well as RNA bases can be obtained. Reducing local beam induced motion on lamellas by Platinum sputter coating results in higher resolution averages while reducing the overall number of required particles. Further investigation can therefore now focus on resolving individual ribosomal states and their arrangement within *e.g*. poly-ribosomes. Finally, this method could bring cellular components, which are less abundant within the reach of high-resolution *in situ* cryo-ET. It thereby opens up new avenues for exploring biology through drug treatment, genetic, and other manipulation directly in living cells.

## Methods

### Sample Preparation

*S. cerevisiae* cells were grown in log phase conditions to an OD_600_ of 0.8. 4 μL of the cells were applied to a glow-discharged 200 mesh holey carbon grid copper grid (Quantifoil R1.2/3) and vitrified in a liquid ethane on a Vitrobot Mark IV (Thermo Scientific) set at 4 °C and 100% humidity. Settings: blot force = 10; blot time = 10 s; wait time = 1 s. Samples were stored under liquid nitrogen until use. Grids were clipped in Autogrids with a cutout slot and subjected to automated FIB-milling on an Aquilos 2 (Thermo Fisher Scientific) using AutoTEM Cryo (Thermo Fisher Scientific) as described elsewhere.^19^ Prior to milling, grids were sputter coated with metallic platinum (Pt) for 30 s using beam current of 30 mA and a pressure of 10 pascal using the in-chamber plasma coater. This was followed by ~ 500 nm coat of organometallic Pt using the gas injection system. After final milling, the lamellas were either used directly for tilt series collection or again sputter coated with a thin layer of metallic Pt. For this post-sputter coating, three different conditions were tested for generating a very thin layer of coat and Pt islands less than 5 nm in size. The final parameter used for high resolution data collection were: beam current: 30 mA, pressure: 10 pascal, voltage: 1 kV, duration: 3 sec.

### Data acquisition

Datasets were collected using a Krios G4 equipped with a Selectris X energy filter and Falcon 4 direct electron detector (Thermo Fisher Scientific). Tilt-series were collected with a dose-symmetric tilt scheme using TEM Tomography 5 software (Thermo Fisher Scientific). The tilt span of ± 60° was used with 3° steps starting at either ± 10° to compensate for the lamella pre-tilt. Target focus was changed for each tilt-series in steps of 0.25 μm over a range of −1.5 μm to −3.5 μm. Data were acquired in EER mode of Falcon 4 with a calibrated physical pixel size of 1.62 Å and a total dose of 3.5 e^−^/Å^2^ per tilt over ten frames. A 10 eV slit was used for the entire data collection. Eucentric height estimation was performed once for each lamella using stage tilt method in TEM Tomography 5 software. Regions of interest were added manually, and positions saved. Tracking and focusing was applied before and after acquisition of each tilt step. The energy filter zero-loss peak was tuned only once before starting the data acquisition.

### Image processing

The data was preprocessed using TOMOgram MANager (TOMOMAN) ^22^. EER images were motion corrected using Relion’s implementation of motioncor^23^. The defocus was estimated using CTFFIND4^24^. Tilt series were aligned using fiducial-less alignment in ARETOMO^25^ Initial tomograms without CTF correction were reconstructed by weighted back projection (WBP) at 16x binning and used for template matching.

Initial particle positions for 80S Ribosomes were determined using the noise correlation template matching approach implemented in STOPGAP ^26^. PDB entry 6gqv^27^ for 80S ribosomes was used to generate a template using the molmap^28^ command in Chimera ^29^. 500 particles per tomogram were picked from 54 and 91 tilt series for +Pt and −Pt datasets, respectively. Subsequent sub tomogram averaging and classification were performed using STOPGAP ^26^. Classification was performed using simulated annealing stochastic hill climbing multi reference alignment as described before ^10^.

Resulting particles for each dataset (~9.5k for −Pt and ~12.5k for +Pt) were then exported to Warp^30^ using TOMOMAN^31^. Subtomograms were reconstructed for Relion 3.0^32^ using Warp at 2x binning (3.2 Å/pix). An iterative approach with subtomogram alignment in Relion and tilt-series refinement in M was performed until no further improvement in gold standard Fourier Shell Correlation (FSC) was obtained. For final averages comparing +Pt and −Pt conditions, ~9.5k particles for each dataset were reconstructed at a pixel size of 2 Å, and another round of subtomogram alignment in Relion and tilt-series refinement in M^21^ was performed until convergence.

For the complete +Pt dataset, particles were further reconstructed at 1x binning (1.6 Å/pix) and a round of subtomogram alignment in Relion and tilt-series refinement in M was performed using a focused mask around LSU.

Densities were visualized and rendered using ChimeraX^33^. In case of the 4.5 Å LSU map for the +Pt dataset, PDB entry 6gqv was docked using rigid body fit in ChimeraX.

## Supporting Information

**Supporting Table 1.**

| **Dataset** | **–Pt** | **+Pt** | **+Pt-extended** |
| --- | --- | --- | --- |
| **Microscope** | FEI Titan Krios G4 | FEI Titan Krios G4 | FEI Titan Krios G4 |
| **Voltage (keV)** | 300 | 300 | 300 |
| **Detector** | Falcon 4 | Falcon 4 | Falcon 4 |
| **Energy-filter** | Selectris X | Selectris X | Selectris X |
| **Slit width (eV)** | 10 | 10 | 10 |
| **Super-resolution Mode** | EER | EER | EER |
| **Å/pixel** | 1.62 | 1.62 | 1.62 |
| **Defocus range (µm)** | -1 to -3.5 | -1 to -3.5 | -1 to -3.5 |
| **Defocus step (µm)** | 0.25 | 0.25 | 0.25 |
| **Acquisition scheme** | -60/60°, 3° | -60/60°, 3° | -60/60°, 3° |
|  | Dose-symmetric | Dose-symmetric | Dose-symmetric |
|  | Tomography 5.0 | Tomography 5.0 | Tomography 5.0 |
| **Total dose (e/ Å<sup>2</sup>)** | ~143.5 | ~143.5 | ~143.5 |
| **Dose rate (e/ Å<sup>2</sup>/sec)** | 7.2 | 7.2 | 7.2 |
| **Tilt series used for STA** | 54 | 64 | 91 |
| **Number of Particles** | 9.5k | 9.5k | 12k |
| **Resolution (Å)** | 6 | 5 | 4.8 (4.5 for LSU) |

**Supplementary Figure 1.**
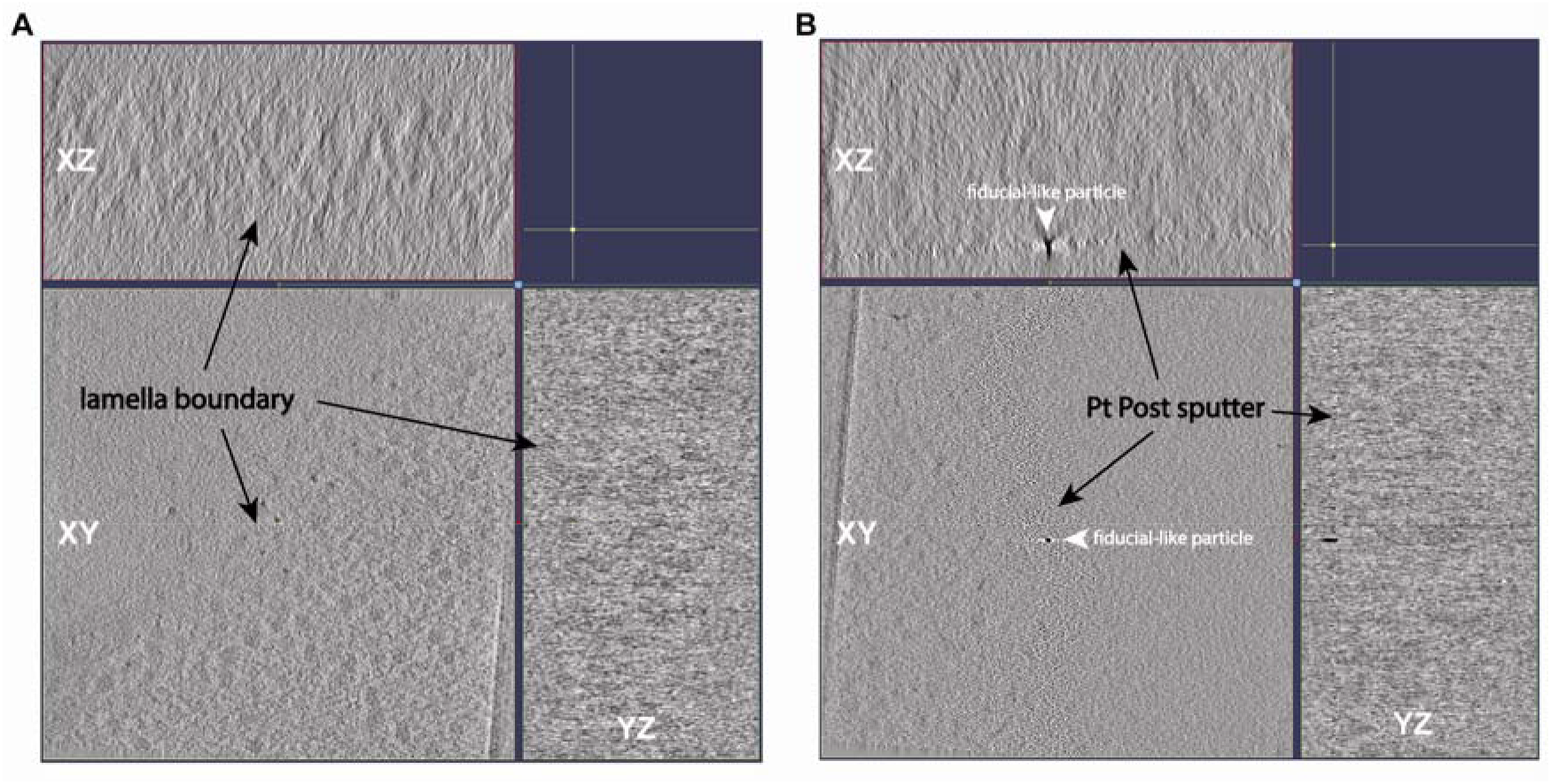
Lamella Coating Optimization. Comparison of A) uncoated and B) ideally coated cryo-FIB lamellas. There is a clear lamella boundary with a finely grained Pt coat. Occasionally, larger fiducial-like particles are produced by the sputter process, which do not negatively affect the tomogram quality.

**Supplementary Figure 2.**
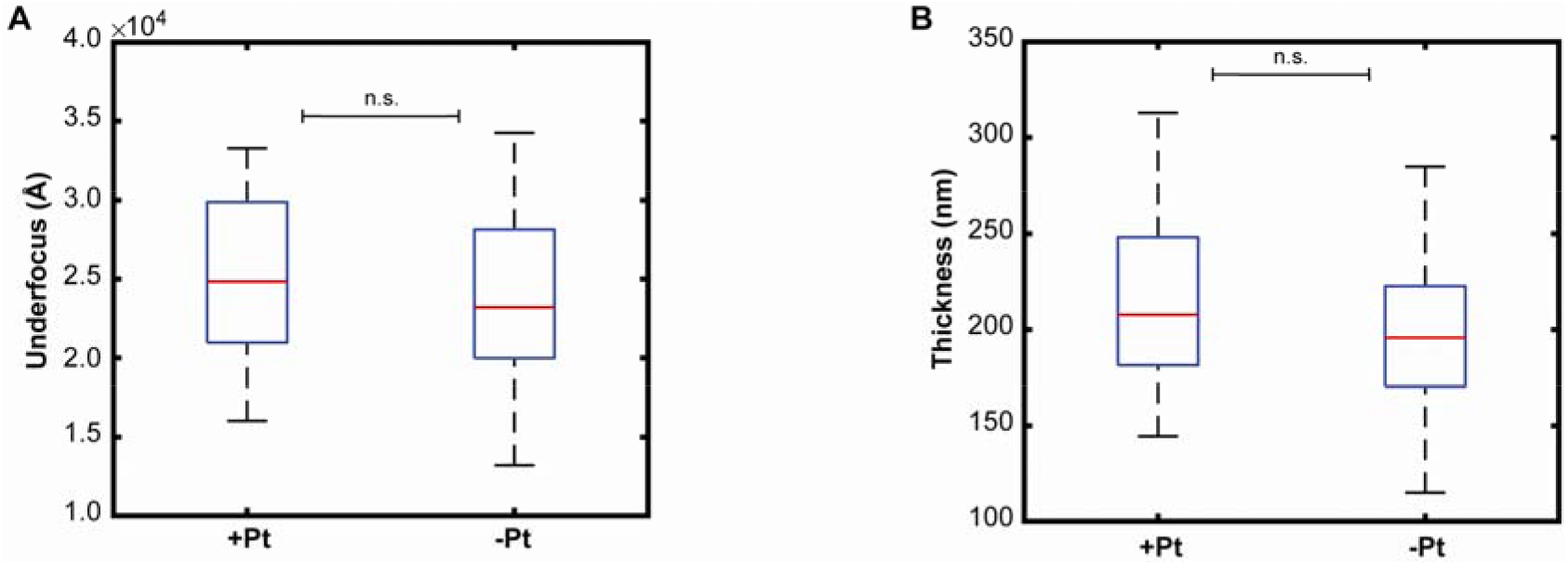
Comparison of Key Lamella Properties. Both A) defocus (underfocus) values and B) lamella thickness are comparable between the +Pt and −Pt datasets. On each box, the central mark indicates the median, and the bottom and top edges of the box indicate the 25th and 75th percentiles, respectively. The whiskers extend to the most extreme data points not considering outliers. Outliers are plotted individually using the ‘+’ marker symbol. P-values are calculated using the two-sample Kolmogorov-Smirnov test. n.s. = not significant.

**Supplementary Figure 3.**
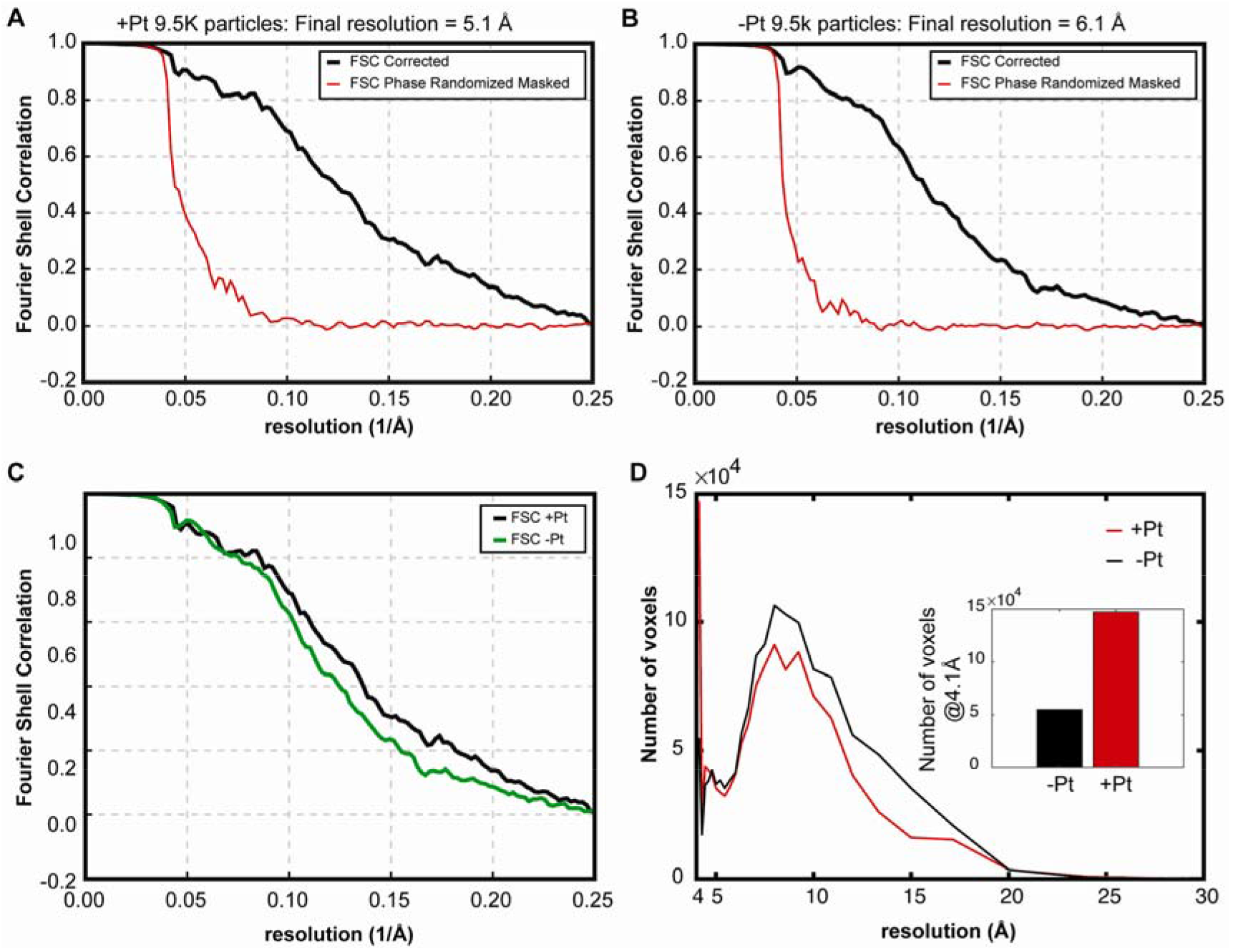
Resolution Potential of the +Pt and −Pt datasets using 9.5k particles each. A) Global resolution of the +Pt dataset at FSC = 0.143 is 5.1 Å. B) Global resolution of the −Pt dataset at FSC = 0.143 is 6.1 Å. C) Overlay of the FSC curves of +Pt and −Pt. D) Histogram of voxels at 4.1 Å for +Pt and −Pt respectively.

**Supplementary Figure 4.**
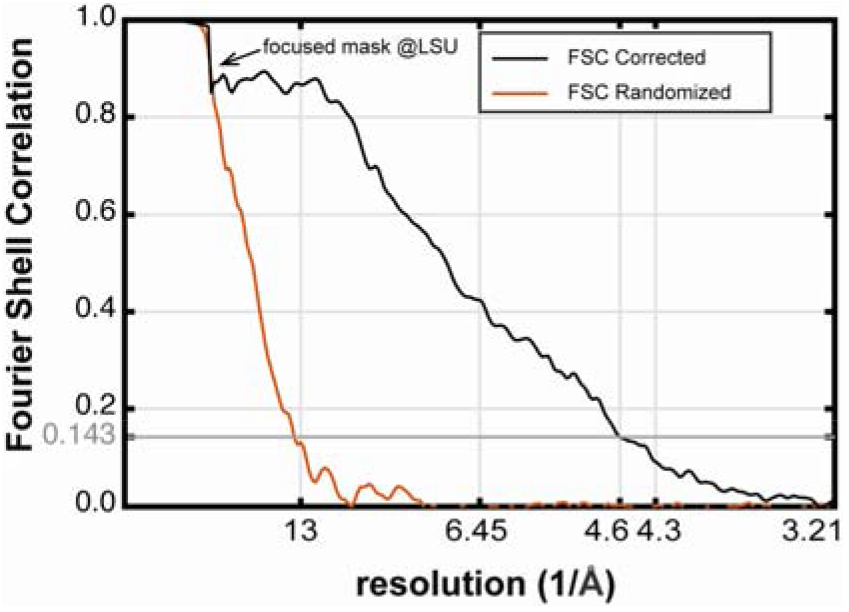
FSC plot of the final + Pt average. With 12.5k particles, the final global resolution of the +Pt dataset is 4.5 Å (0.143 FSC cutoff).

